# Failures that are not failures: unobserved sensory history predicts escape behavior

**DOI:** 10.64898/2026.07.17.739074

**Authors:** Pranay Dasyam, Joby Joseph

## Abstract

Animals often fail to respond to reliable sensory stimuli, a phenomenon commonly attributed to stochastic neural noise or modulation by internal states. In locusts, looming-evoked jump escape response (JER) fails in ~40% of trials, and these failures correlate with multiplexed features of the descending contralateral movement detector’s (DCMD) response^1,2^. Using *Hieroglyphus banian*, we show that these failures are predictable from an unobserved “Light-to-Dark transition” (LDT) caused by the procedure in the assay, minutes before the looming stimulus. In the JER assay, LDT induced a long-lasting competing behavioral state, grooming, which suppressed JER. Reducing the amplitude of this transition by illuminating the tunnel decreased grooming and rescued JER. Grooming was reliably induced by LDT in the grasshopper under culture conditions. Electro-mechanical induction of grooming was sufficient to cause JER failure. The grooming state induced by either LDT or electro-mechanical stimulation reduced looming-evoked DCMD responses through a divisive gain modulation while preserving normalized variability. Grooming was rare under natural field conditions. Thus, accounting for previously unobserved stimulus history can explain a substantial component of apparent behavioral failures that would otherwise be attributed to stochastic neural noise.

**Highlights:**

- An unobserved prior sensory event predicts escape failures.
- Light-to-Dark Transition (LDT) induces a long-lasting grooming state.
- Grooming state suppresses escape responses via divisive gain reduction in the DCMD neuron.
- Behavioral failures can arise from sensory history rather than neuronal noise.

**Graphical abstract:** 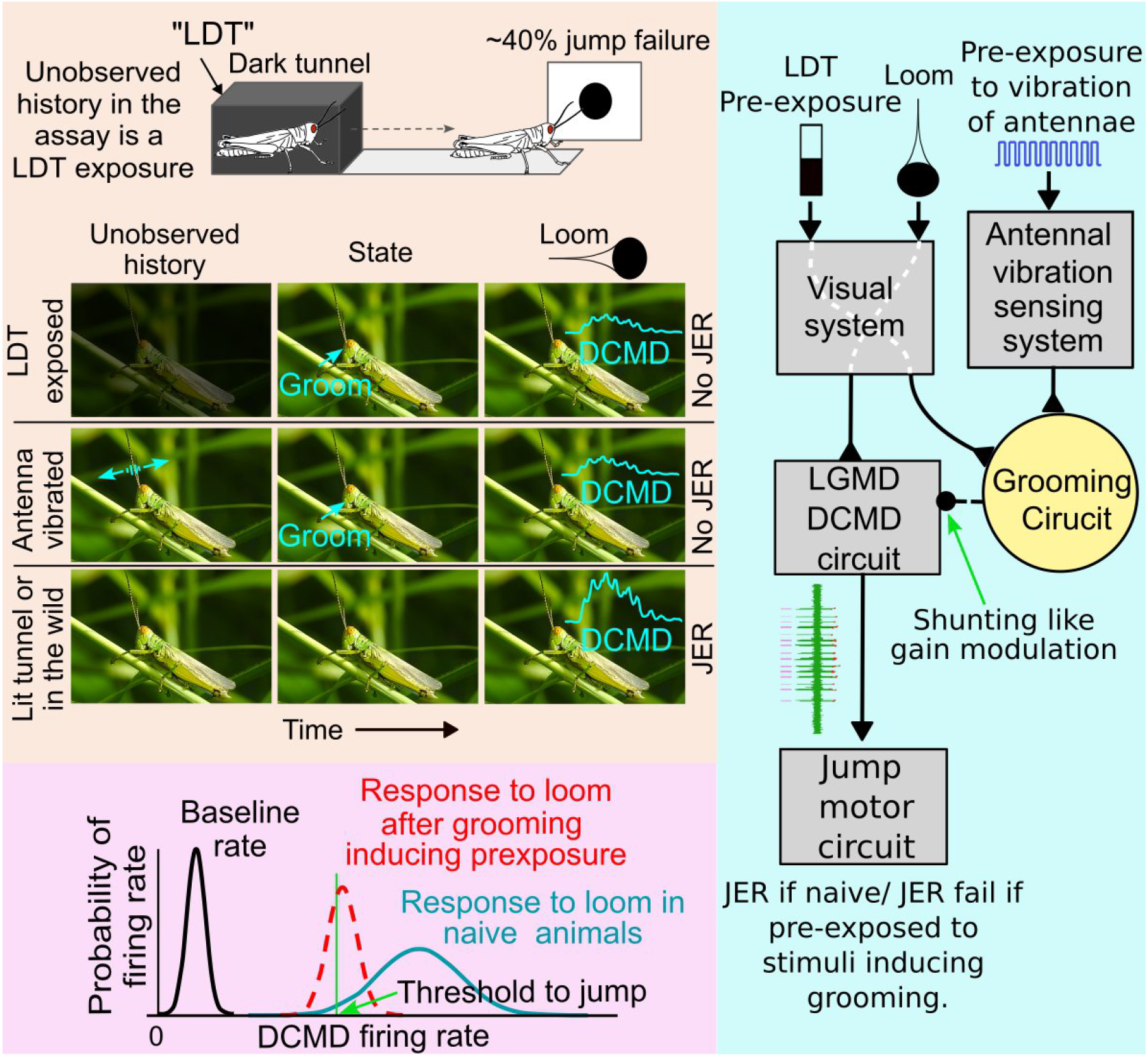

## Introduction

Behavioral responses to deterministic sensory stimuli often exhibit substantial trial-to-trial variability, commonly attributed to stochastic noise in neural processing ^*3–5*^. However, this variability may instead reflect modulation by internal states^*6–8*^. These internal states may in turn be deterministically caused by unobserved stimulus history. Internal states such as satiation or stress can modulate behavioral responses, and competing motor programs may influence decision-making ^*9–12*^. Distinguishing between stochastic and structured sources of intra-individual variability ^*13*^ remains a central challenge in understanding how neural circuits generate reliable behavior. Escape response assays are useful for studying decision-making, as these behaviors are rapid, survival-critical, and mediated by relatively well-defined neural circuits. Despite being survival-critical, escape behaviors fail at substantial rates across species, including insects, crustaceans, and vertebrates ^*9,14–20*^. In grasshoppers and locusts, looming-evoked escape is mediated by the lobula giant movement detector-descending contralateral movement detector (LGMD–DCMD) pathway, which provides a direct link between sensory input and motor output ^*14,21,22*^. Although this behavior is critical for survival, JER failures are high in these assays, independent of the angular speed (l/|v| ratio) (Figure 1A) ^*21,23–26*^. This behavioral assay and the underlying neural circuit have been used extensively to gain insight into fundamental principles of neural computation, like multiplication operation and variability in decision-making in neural circuits ^2,26,27^. Two of the three multiplexed parameters in the DCMD response to the loom have been shown to be correlated to the success of the jump^2^. However, these results do not explain why there is such a high rate of failure in the JER, a survival-critical behavior. Starting from the serendipitous observation that animals often groom during trials in which JER fails, we show that a substantial fraction of looming-evoked JER failures can be explained by a long-lasting grooming state induced by an unobserved prior event, a light-to-dark transition (LDT). We show that this grooming state reduces the DCMD responses by divisive gain modulation. We could rescue the success rate in the JER assay by reducing the amplitude of the light-to-dark transition. In the natural field conditions, *Hieroglyphus banian* shows little grooming, implying a likely high success rate of JER in the field. These findings reveal that trial-to-trial behavioral variability can originate from a hidden sensory history that induces structured modulation of the internal state rather than stochastic noise in neural circuits.

**Figure 1.**
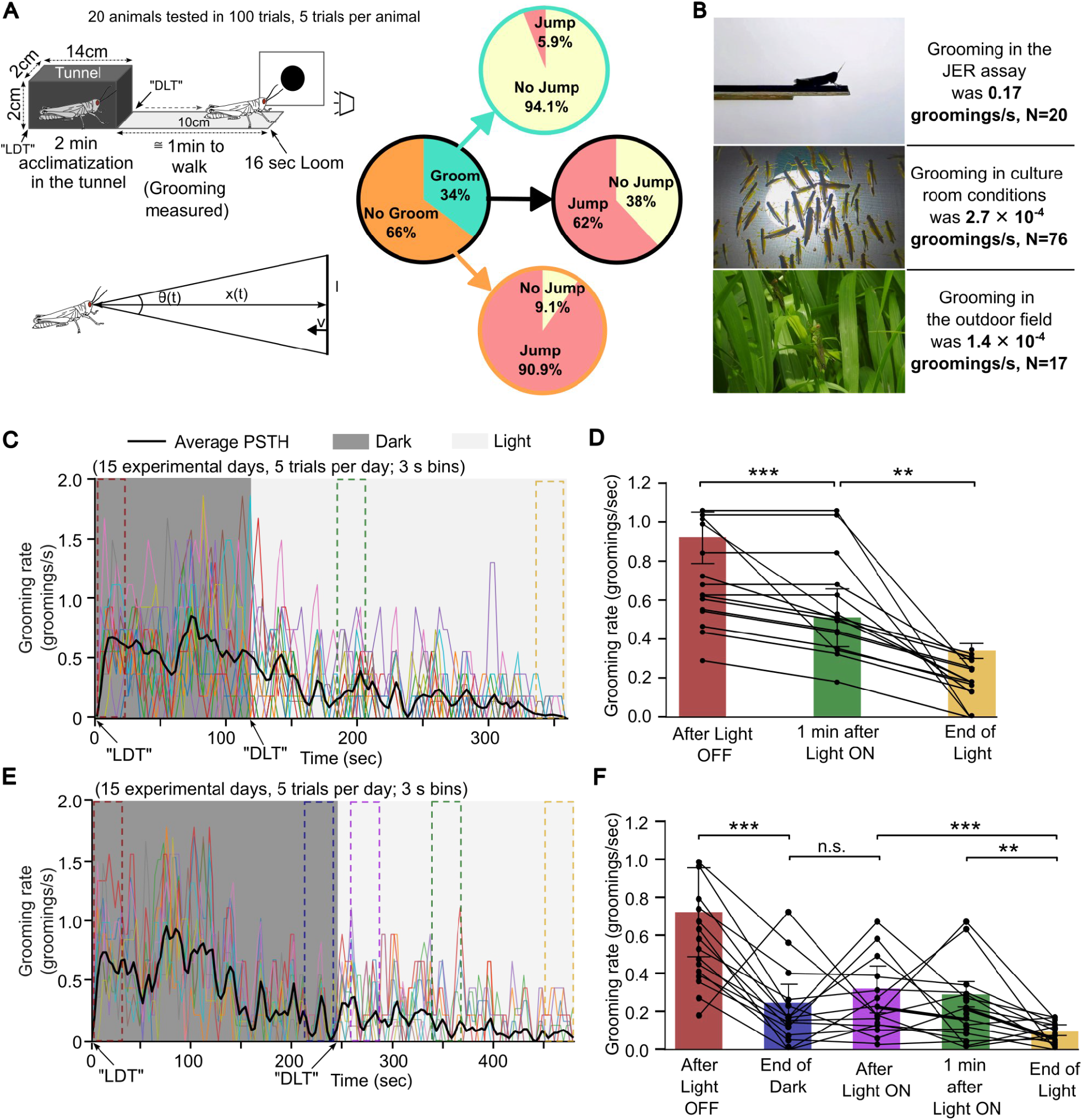
LDT in the JER assay induces grooming, and grooming rate correlates with JER failure. **(A)** Left: Schematic of the JER assay and looming stimuli parameters. Right: Correlation between grooming and JER in *Hieroglyphus banian*. Across 100 trials (N = 20 animals, 5 trials/animal), grooming occurred in 34%, and JER failed in 38% of trials. JER failed in 94.1% of grooming trials compared to 9.1% of non-grooming trials (GEE, β = −5.21, OR = 0.0054, p = 4.8*10^-9^). Illumination intensity was 170 lux outside the tunnel and 0 lux inside (“Dark”). **(B)** Grooming rates in the JER assay were very high compared to natural environments. **(C)** Peristimulus time histogram (PSTH) of grooming in the culture chamber following a pre-exposure to LDT under conditions analogous to the JER assay. The first 120 s corresponds to “Dark” followed by a prolonged (240 s) “Light” condition. Dashed boxes indicate the time windows used for quantification in (D). In (C) and (E), colored traces represent individual days; the black trace shows the mean PSTH smoothed by a 4-bin-wide moving average, and dashed boxes indicate time windows used for quantification. **(D)** Grooming rate (mean rate within dashed windows in (C), ± SEM) decreased significantly from “After Light OFF” to “1 min after Light ON” (p = 0.0008) and further to “End of Light” (p = 0.006) (one-way ANOVA with Tukey’s post hoc test). In (D) and (F), points represent averages per day (5 trials), with lines connecting the same days. **(E)** PSTH of grooming during cyclic “Light” and “Dark” of equal durations. The first 240 s corresponds to “Dark”, followed by 240 s of “Light”, though the stimulus was cyclic, and the starting phase was randomized among days. **(F)** Grooming rate (mean rate within dashed windows in (E), ± SEM) across five time windows indicated in (E). Grooming decreased from “After Light OFF” to “End of Dark” (p = 0.0004), showed no significant difference between “End of Dark” and “After Light ON”, and decreased from “After Light ON” to “1 min after Light ON” (p = 0.0006) and to “End of Light” (p = 0.0016) (one-way ANOVA with Tukey’s post hoc test). JER: jump escape response, LDT: Light-to-dark transition, DLT: Dark-to-light transition.

## Results

### Grooming behavior is correlated with failure of looming-evoked jump escape responses

To investigate the relationship between grooming behavior and looming-evoked JER in *Hieroglyphus banian*, we quantified grooming and escape responses in 20 animals across 100 trials (5 trials per animal). Animals engaged in grooming in 34% of trials (Figure 1A, left), whereas JER failed in 38% of trials (Figure 1A, middle). JER failed in 94.1% of the grooming trials (Figure 1A, top). In contrast, JER was successful in 90.9% of the non-grooming trials (Figure 1A, bottom). Thus, grooming is highly correlated with JER failure. These results demonstrate a strong behavioral state-dependent modulation of the probability of jump escape response.

### Light-to-dark transition induces a long-lasting grooming state

Because grooming strongly predicted JER failure, and grooming rate was high in the assay, we measured the spontaneous grooming rate of *H. banian* in the culture chamber and in the field. Grooming rates were 2.7 × 10^-4^ s^-1^ in the culture chamber and 1.4 × 10^-4^ s^-1^ in the outdoor field recordings (Figure 1B). Surprisingly, these grooming rates were nearly zero compared to that in the JER assay (0.17 s^-1^), indicating that grooming in the assay was not spontaneous but induced by some aspect of the experimental setup. In the JER assay, to motivate them to walk to the point of the loom presentation, animals were initially placed in a dark tunnel, causing an LDT, before opening the tunnel door onto an illuminated platform. We therefore tested whether grooming is triggered in the culture chamber by LDT, corresponding to the introduction into the dark tunnel from the bright ambient condition (see Materials and Methods). Following a prolonged ambient “Light” condition, switching to the “Dark” condition resulted in a marked increase in grooming rate, which decreased slowly over time (Figure 1C). Grooming rate measured immediately after LDT was significantly higher than grooming rate measured “1 min after return to the Light ON” condition (p = 0.0008). Grooming rate further decreased from “1 min after Light ON” condition to the “End of Light” (p = 0.006) (Figure 1D). These results demonstrate that LDT robustly increases grooming behavior, and the elevated grooming persists for minutes even after the Light is turned ON. The grooming rate at the time point corresponding to the looming stimuli in the JER assay (“1 min after Light ON”, green) is of the order of 10^4^ times higher than the spontaneous rate in the culture chamber and the outside field. Thus, a previously unobserved sensory stimulus, the LDT, preceding the looming stimulus, accounts for increase in grooming in the JER assay.

### LDT, but not dark-to-light transitions (DLT), induce grooming

Asymmetry in the grooming response induced by LDT compared to DLT may indicate ecological significance. To determine whether grooming is triggered specifically by an LDT or by light transitions in general, we quantified grooming under cyclic “Light” and “Dark” conditions of equal duration, with the starting phase balanced across days (Figure 1E). LDT, but not DLT, increased grooming rate (Figure 1E and 1F). Grooming increased abruptly at the LDT, then gradually decreased during the “Dark” phase and continued to decline during the subsequent “Light” phase (Figure 1E and 1F). Together, these results demonstrate that grooming is specifically initiated by LDT rather than by light transitions in general, indicating the LDT as the sensory event responsible for the persistent grooming observed in JER assay.

### Illuminating the tunnel reduces grooming and rescues JER

If the LDT in the tunnel induces grooming and causes failure of looming-evoked JERs, then illuminating the tunnel, as in the natural habitat, should reduce grooming and rescue JER. In dark tunnel (0 lux) trials, grooming occurred in 37.1%, and JER failed in 48.9%. In the tunnel-illuminated (98 lux, rescue) trials, grooming decreased to 4% of trials, and overall JER failure decreased to 20%. Thus, tracking the occurrence of the LDT alone was sufficient to predict JER failure, even without directly monitoring grooming behavior. In the dark tunnel condition, JER failed in 88.3% of the grooming trials and 25% of the non-grooming trials. In the illuminated tunnel condition, JER failed in 100% of the grooming trials and 16.3% of non-grooming trials. Overall, grooming was significantly reduced, and JER success increased in the illuminated tunnel condition compared to the dark tunnel condition (Figure 2A). Grooming was correlated with JER failure in both conditions, Illuminated: GEE, β = −24.20, OR = 0, p = 0) and Dark (GEE, β = −3.11, OR = 0.0444, p = 0.000425). Together, these results indicate that tunnel illumination increases jump escape success primarily by reducing the incidence of grooming, and that under natural field conditions, JER success for *H. banian* is high. More importantly, accounting for this previously unobserved sensory event substantially improved prediction of JER failures, even without directly monitoring grooming behavior.

**Figure 2.**
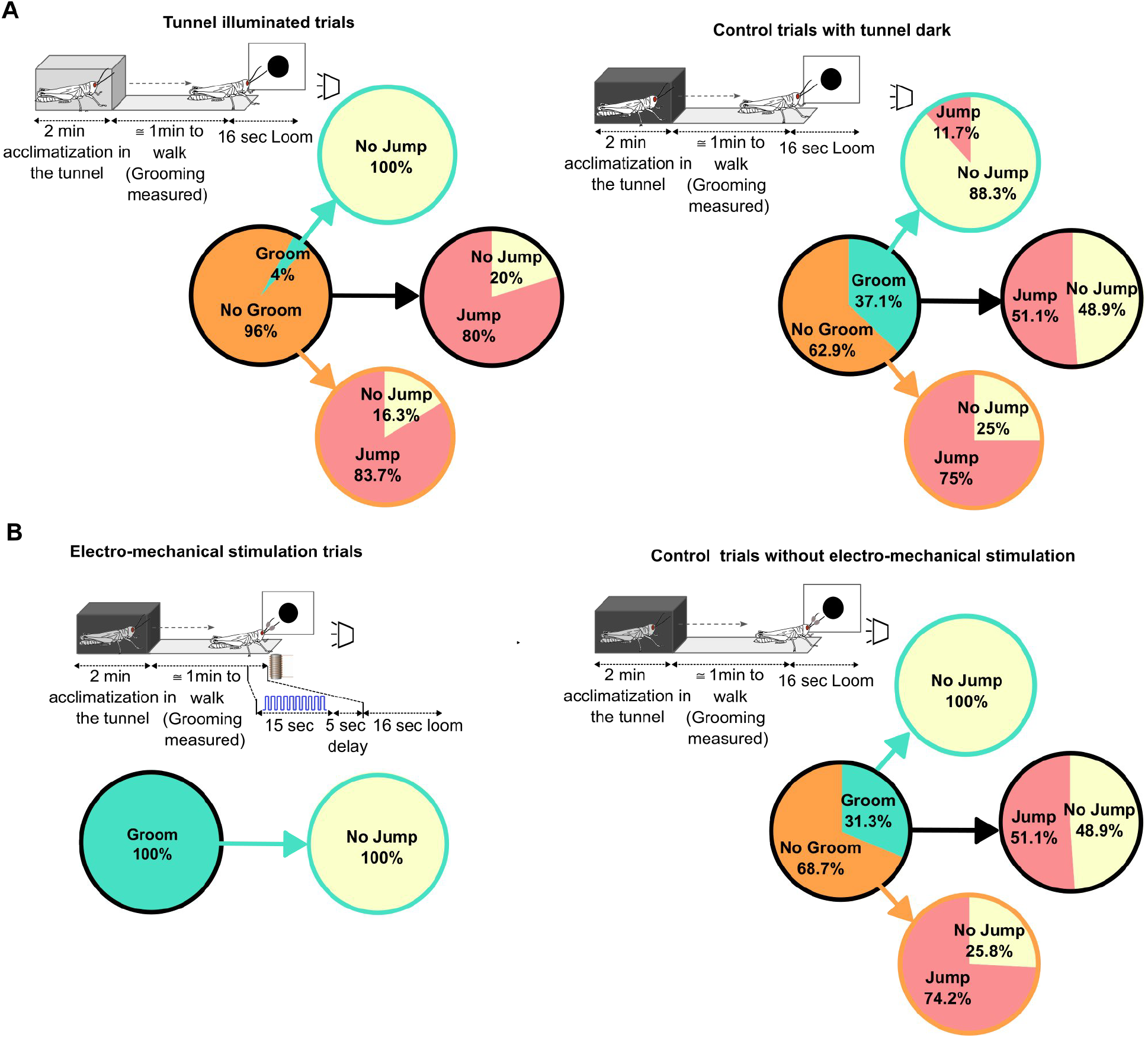
Illuminating the tunnel rescues JER, and electro-mechanical induction of grooming eliminates JER. **(A)** Grooming rate and JER success rate when the tunnel was illuminated (98 lux, left) vs when the tunnel was dark (0 lux, right) for the same grasshoppers. Grooming rate decreased (GEE, β = −2.57, OR = 0.08, p = 0.00574), and JER increased (GEE, β = 1.34, OR = 3.83, p = 6.57*10^-05^) with the tunnel illuminated. **(B)** Grooming rate and JER success rate with electro-mechanical stimulation of antennae (left) vs no electro-mechanical stimulation (right) for the same grasshoppers. Grooming rate increased to 100% (GEE, β = 26.36, OR = 2.8*10^11^, p = 0), and JER decreased (GEE, β = −25.61, OR = 0, p = 0.0057). For (A) and (B), N = 15 grasshoppers, with three trials for each condition. Treatments and controls were alternated and counterbalanced between animals.

### Electro-mechanically induced grooming was sufficient to suppress looming-evoked JER

To test whether grooming itself is sufficient to suppress looming-evoked JER, we electro-mechanically induced grooming by stimulating the antennae before presentation of looming stimuli and compared behavioral outcomes with control trials without electro-mechanical stimulation. During electro-mechanical stimulation trials, animals received 10 pulses of 15 s duration (50% duty cycle; 0.75 s ON, 0.75 s OFF; 0.67 Hz), which reliably evoked grooming in all trials (Figure 2B). Grooming occurred in 100% of stimulated trials and resulted in 100% failure of JER. In contrast, grooming occurred in 31.3%, and JER failed in 48.9% of the control trials. JER failed in 100% of grooming trials and 25.8% of non-grooming trials. Thus, grooming was correlated with JER failure in both conditions, with electro-mechanical stimulation (GEE, β = −12.78, OR = 0, p = 0) and without electro-mechanical stimulation (GEE, β = −25.62, OR = 0, p = 0). Together, these results indicate that grooming, or the underlying cause of grooming, is sufficient to suppress looming-evoked jump escape responses, demonstrating that the behavioral failures can be recreated by inducing the same internal state.

### Pre-exposure to LDT reduces response gain and preserves normalized variability in DCMD

Having established that LDT increases grooming and that grooming is associated with failure of JER, we tested whether prior exposure to this transition alters neural activity in the DCMD, whose firing rate predicts jump success ^2^. Looming-evoked responses of the DCMD were recorded extracellularly from the ventral nerve cord using hook electrodes (see methods, Figure S1) following pre-exposure to either “Light” or “LDT” (Figure 3A and 3B). The two pre-exposure conditions were randomly alternated within each animal. Baseline firing rates did not differ between the two pre-exposure conditions (Figure 3Bi). However, the looming-evoked DCMD firing rate was significantly lower following pre-exposure to LDT compared to “Light” condition. (Figure 3B). Mean firing rate within a window around the peak was significantly reduced when the animal was pre-exposed to LDT (Figure 3B ii and 3Biii).

**Figure 3.**
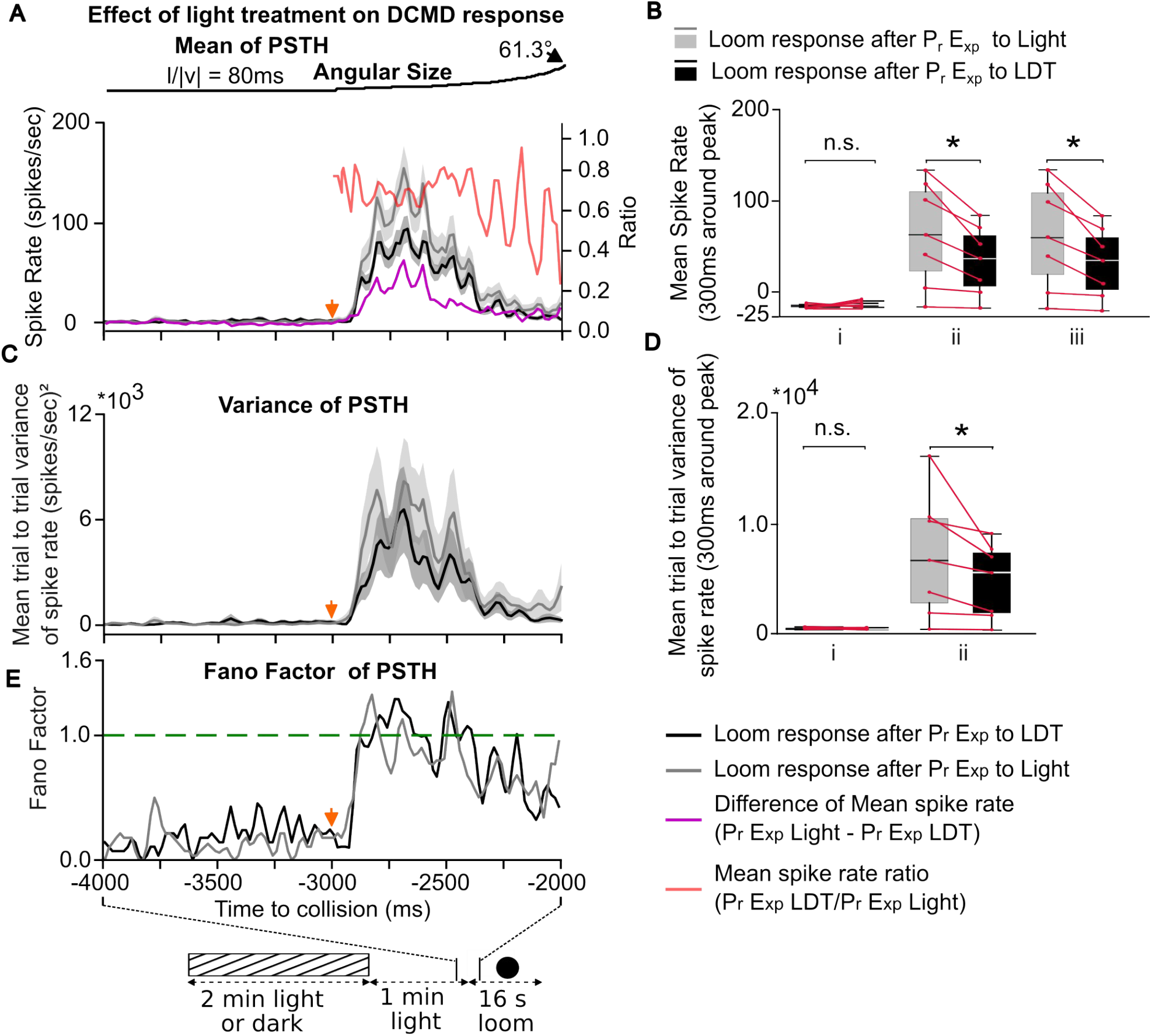
Pre-exposure to LDT reduces DCMD firing rate, variance, and gain in *Hieroglyphus banian*. **(A)** Mean DCMD firing rate ± SEM (PSTH bin width = 1/60 s, N = 7, 5–10 trials per animal) aligned to looming stimuli onset (orange arrow) following pre-exposure to “Light” (gray) or LDT (black). Difference (magenta) and mean of the ratio of spike rate (red; right axis) between conditions (“Light” – “LDT”; “LDT” / “Light”). **(B)** Paired comparisons of firing rate across animals (300 ms window around peak response): (i) baseline rate, (ii) looming-evoked rate (p = 0.014), and (iii) looming-evoked rate above baseline (p = 0.0164). Each point represents the trial-averaged response of an individual animal, with lines connecting paired conditions. **(C)** Trial-to-trial variance of DCMD firing rate (mean across animals) for “Light” (gray) and “LDT” (black) pre-exposure conditions. **(D)** Paired comparisons of variance across the same time windows as in (B): (i) baseline and (ii) looming-evoked variance (p = 0.021). **(E)** Fano factor (variance/mean) of DCMD firing rate for both conditions. The green line indicates Fano factor = 1. All PSTHs were smoothed using a rectangular moving-average filter (window length = 4 bins) and all tests were paired *t*-tests.

The pointwise difference and mean of the ratio of spike rates between conditions are shown in Figure 3A (magenta and red traces, respectively). The difference trace followed a temporal profile similar to the looming-evoked response, whereas the ratio of spike rates (“LDT” / “Light”) remained approximately constant across the evoked response window, consistent with a multiplicative scaling of the looming-evoked response (Figure S6).

Trial-to-trial variance of the looming-evoked spike rate was also reduced following LDT pre-exposure compared to “Light” (Figure 3C and 3D; p = 0.021). The Fano factor, which is a measure of variability relative to the mean, remained close to 1 during the looming response in both conditions (Figure 3E), consistent with what is reported in cortical cells having gain modulation via shunting inhibition ^28^. Thus, a reduction in DCMD response while maintaining a constant Fano factor can provide a neural correlate for the increased failure rate of JER following introduction into the dark tunnel in the JER assay, demonstrating that a previously unobserved sensory event can produce persistent neural modulation that explains a substantial fraction of the behavioral failures.

### Pre-exposure to electro-mechanical stimulation of the antennae before the looming stimuli reduces response gain and preserves normalized variability in DCMD

Given that electro-mechanically induced grooming suppresses jump escape responses (Figure 2B), we tested whether pre-exposure to electro-mechanical stimulation reduces looming-evoked DCMD activity (Figure 4A). The two pre-exposure conditions were alternated within each animal. Baseline firing rates did not differ between the two conditions (Figure 4A). However, looming-evoked DCMD firing rate and variance were significantly reduced following electro-mechanical pre-exposure (Figure 4B and 4D). Electro-mechanical stimulation produced effects similar to those observed following LDT pre-exposure (compare Figure 3 and Figure 4). Together, these results indicate that pre-exposure to electro-mechanical stimulation reduces the gain of the DCMD pathway while preserving the proportional relationship between mean firing rate and variance (Figure S7). This reduction in neural response provides a physiological correlate for the suppression of jump escape responses observed following grooming-inducing electro-mechanical stimulation.

**Figure 4.**
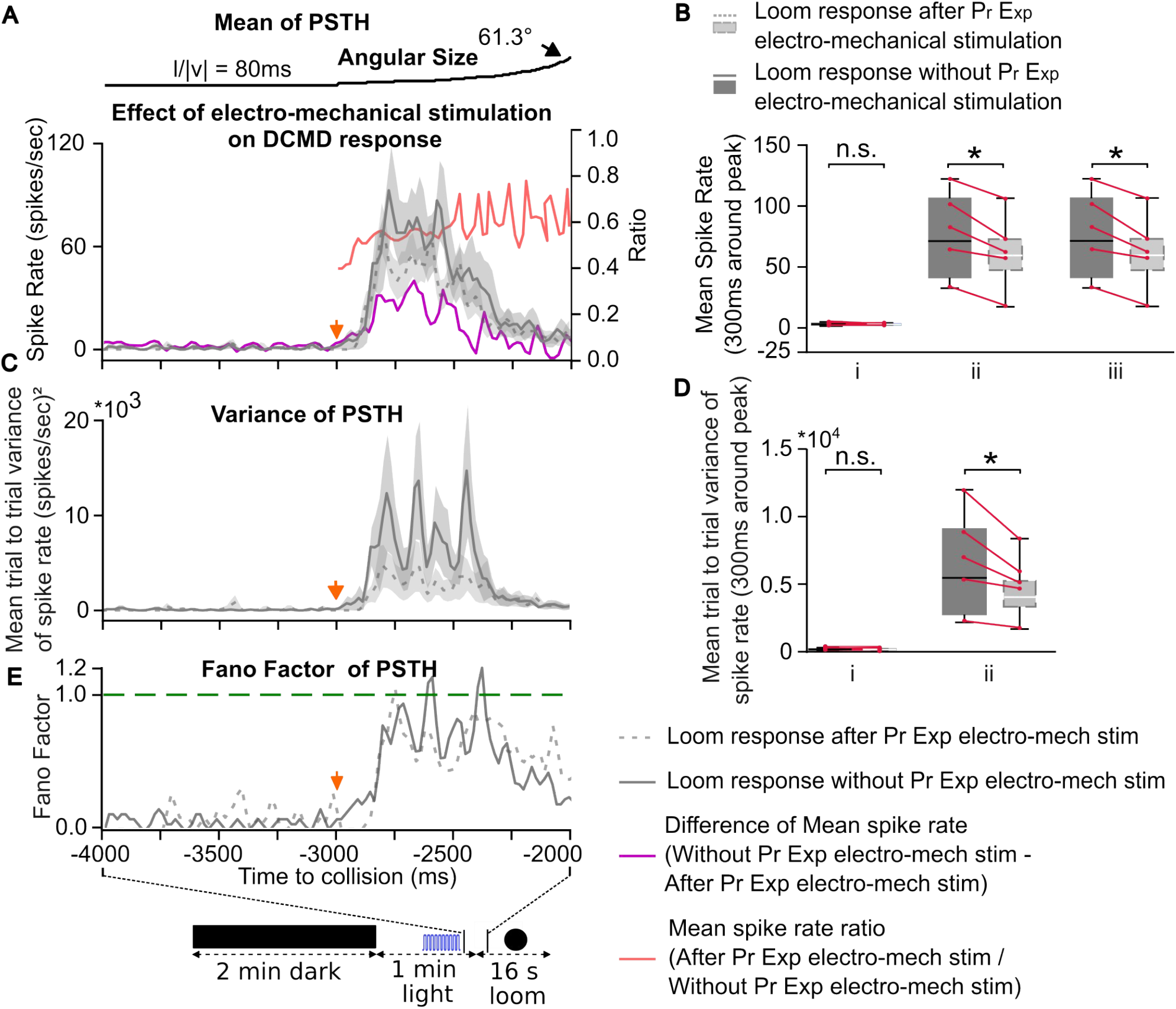
Pre-exposure to electro-mechanical stimulation of antennae before the looming stimuli reduces looming-evoked DCMD firing rate, variability, and gain in *H. banian*. **(A)** Mean DCMD firing rate ± SEM (PSTH bin width = 1/60 s; across 5 animals) aligned to looming stimulus onset (orange arrow), with (dotted gray) and without (solid gray) electro-mechanical pre-exposure. Difference (magenta) and mean spike rate (red; right axis) between the two conditions. **(B)** Paired comparisons of firing rate across animals (300 ms window around peak response): (i) baseline, (ii) looming-evoked rate (p = 0.032), and (iii) looming-evoked rate above baseline (p = 0.037). Each point represents one animal (trial-averaged), with lines connecting paired conditions. **(C)** Trial-to-trial variance of DCMD firing rate (mean across animals) with (dotted gray) and without (solid gray) electro-mechanical pre-exposure. **(D)** Paired comparisons of variance across the same windows as in (B): (i) baseline and (ii) looming-evoked variance (p = 0.041). **(E)** Fano factor (variance/mean) of DCMD firing rate for both conditions. The green line indicates Fano factor = 1. All PSTHs were smoothed using a rectangular moving-average filter (window length = 4 bins).

## Discussion

### Behavioral failures reflect hidden internal state rather than stochastic noise

We showed that a substantial fraction of apparent failures in the JER are not random but are associated with an alternative behavioral state, grooming, induced by a prior, unobserved cause. Thus, failure of escape behavior can arise from structured, internal state-dependent modulation of neural activity rather than stochastic neuronal noise. Light transitions (LDT and DLT) elicited a response in the DCMD neuron only at the transition (Figure S4). The electro-mechanical stimulation did not cause any immediate response in the DCMD spike rate at all (Figure S5). Thus, neither the JER failure nor the grooming was dependent on direct activation of DCMD by these stimuli, indicating that the effects arise from a persistent internal state induced by prior sensory experience.

DCMD firing rate in response to looming stimuli is a strong predictor of escape response ^2,21,23,24,26,29–31^. We showed that grooming-inducing conditions reduce the gain of the DCMD pathway. This reduction in the gain is consistent with increased JER failure, as DCMD firing rate failed to reach a threshold in the time required to initiate co-contraction and jump before collision. It is also interesting that this grooming state might have been accidentally induced by the experimental setup in ^2^ and only that enabled them to probe the parameters of DCMD response to looming stimuli that correlate with the success of JER.

There is very little grooming in the wild in *H. banian*. Therefore, DCMD responses to looming stimuli in the wild are likely to have a higher mean and variance than in the assay. This would then imply a higher jump escape success, as expected for a survival-critical behavior. Other descending neurons are also shown to be involved in JER ^12,32,33^ but we have not examined them in this study.

### Light transition induced grooming: a novel stimulus-induced state

To our knowledge, this is the first report of grooming induced by light transitions in any species. In insects, grooming has been extensively studied as a context-dependent behavior ^9,33^, including in cockroaches after a wasp sting (venom) ^35^. How grooming circuits interact with escape circuits remains unknown. Dopamine is involved in inducing grooming in cockroaches caused by a wasp sting ^35^. It is therefore possible that transitions from light to dark induce stress and subsequent dopamine release, leading to grooming in *H. banian* and modulating DCMD firing rate. DCMD firing rate is reduced under grooming conditions by a constant scaling factor while maintaining a near constant Fano-factor, consistent with shunting inhibition reported in some systems (Figure S6 and S7) ^36,37^. Abrupt transitions from light to dark occur in natural environments only when eyes are obstructed, and grooming may be a mechanism to clear the obstruction. It is tempting to view this ongoing grooming as a task the grasshopper is engaged in, and thus the failure to evoke JER is due to a task-switching cost. The interaction between grooming and escape circuits also raises the possibility that proprioceptive feedback associated with grooming can contribute to the observed modulation, where competing motor programs are coordinated through mutual inhibition or hierarchical control ^38,39^.

### Unobserved sensory history improves behavioral prediction

These findings have important implications for how variability in neural systems is interpreted. Though neuronal noise could always be present and play a role in the exploration-exploitation trade-off, the cause of failure of behavioral response that is often attributed to “noise” may instead reflect structured, state-dependent dynamics arising from deterministic sensory events in the recent past. In this particular case, the LDT-induced internal state resulted in an external manifestation, grooming. Such observable external manifestation of internal state may not be present for all stimulus events in the animal’s past. Importantly, our ability to predict JER failure did not depend on observing grooming itself. Thus, even when such manifested behavior is not evident, tracking the stimulus history can improve behavioral prediction, as demonstrated by the improved prediction of JER failure using LDT history (Figure 2). Thus, in this study, accounting for a previously unobserved stimulus history substantially explained behavioral failures that would otherwise appear to be caused by internal noise. More generally, our results support the idea that behavioral prediction may be improved by tracking more of the organism’s prior environment history, even when there is no obvious immediate observable response to those events.

## Resource availability

### Lead Contact

Requests for further information and resources should be directed to and will be fulfilled by the lead contact, Joby Joseph.

### Data and code availability

The data that support the findings of this study are available from the lead author upon reasonable request.

## Author contributions

Conceptualization: P.D., J.J.; Formal analysis: P.D., J.J.; Investigation: P.D.; Methodology: P.D.,J.J.; Supervision: J.J.; Visualization: P.D., J.J.; Writing – original draft: P.D.; Writing – review & editing: P.D., J.J. **Other**: Setup: P.D., Jing Tazong Taruk,

## Supplementary Information

Document S1: Supplementary Figures S1 to S7 and their legends.

Movie S1: Representative looming-evoked jump escape response.

Movie S2: Representative grooming behavior following a light-to-dark transition (LDT).

## Experimental Model

### Grasshopper (*Hieroglyphus banian)*

Adult male and female *Hieroglyphus banian* were used. Animals were reared at the Centre for Neural and Cognitive Sciences, University of Hyderabad, India, under a 14 h light / 10 h dark cycle at 25 to 32°C and 60 to 95% relative humidity. Light intensity in the colony was ~25,700 lux.

## Methods details

### Visual stimuli and behavioral assay

Looming stimuli were generated using custom Python scripts (OpenCV) and consisted of a dark disc expanding on a bright background. The angular size of the stimulus was *θ*(*t*) = 2*tan*^−1^(*l*/*vt*), where *l* represents half the object size, *v* the approach velocity, and *t* time to collision ^26^. Stimuli were presented at a *l*/|*v*| ratio of 80 ms (50 ms in a separate condition; Figure S3) on a tablet screen (Lenovo P53) (25.2 × 15.9 cm) positioned 6.5 cm from the animal’s eye. Behavioral assays were conducted in a 3D-printed apparatus adapted from a previous study ^26^. A frontal light exposure (170 lux) induced phototactic walking in the grasshopper. Tunnel illumination was either 0 lux (dark condition) or 98 lux (illuminated condition), measured directly using a digital lux meter (LX-101A, HTC) or using a photodiode (BPW21R) calibrated with the lux meter in the case of a tunnel. Illumination intensity of the tunnel was determined to the brightest value at which the animal would still show phototactic walking towards the platform. For each trial, animals were placed in the tunnel for 2 min, after which the front door was opened to allow movement onto the platform. Looming stimuli were presented when the animal reached the end of the platform, such that one eye was approximately aligned with the center of the display. Trials in which animals failed to reach the end position were excluded. Each animal was subjected to five trials with a 10-minute inter-trial interval. Jump Escape Response (JER) probability was defined as the fraction of trials in which a jump occurred following a looming stimulus presentation.

### Grooming quantification and experiment manipulations

Grooming behavior (antennal, eye, and mouthpart grooming) was manually scored from video recordings. Grooming rate was defined as grooming events per second calculated by dividing the total time for which the animal groomed by the time duration of observation. For culture chamber and field recordings, grooming bouts were counted across the entire observation period and normalized by the number of animals present within the field of view to obtain grooming events/s.

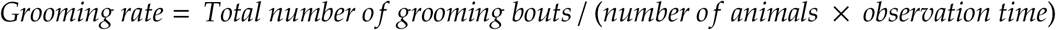

In the JER assay, grooming was quantified for a single animal per trial, and grooming bouts were counted during the interval from tunnel exit until the onset of the looming stimulus. Grooming rates were therefore normalized by the duration of this interval. Grooming rate (count/s) was calculated in predefined 30 s time windows. For each experimental day, grooming rates were averaged across five trials to obtain a single data point per day, which was used for box-and-whisker plots and statistical comparisons. To determine baseline grooming levels under culture conditions, spontaneous grooming behavior was recorded in the culture chamber using an infrared (IR) wildlife camera (64 MP resolution, 1080p video, night vision enabled; recordings monitored remotely using Game Camera Pro 2). A double convex lens (5 diopters; focal length 20 cm; diameter 50 mm) was mounted in front of the camera to obtain focus at short distances. Recordings were performed for approximately 1 h per day over 5 consecutive days, with 20-25 animals within the field of view per day. To assess grooming behavior under natural conditions, field recordings of *H. banian* were obtained in grass patches using a Nikon Z camera equipped with a Nikon 100-600 mm lens. Approximately 1 h of video per day was recorded across 12 days. Grooming events were subsequently quantified from video recordings using the same criteria described above.

Light transition assays were performed using controlled illumination sequences: (i) 2 min Dark followed by 4 min Light, and (ii) 4 min Light followed by 4 min Dark. Five trials per day/animal were conducted for 15 days, and grooming rates were quantified in predefined time windows.

To test the effect of illumination, animals were tested alternately in a dark tunnel (0 lux) and an illuminated tunnel (98 lux). If the tunnel was equally lit as the light intensity outside, the grasshopper would not walk towards the external light. This 98lux was arrived at by finding the highest illumination inside the tunnel at which the grasshopper would still walk towards the external light. Each animal underwent 45 trials per condition with a 10 min inter-trial interval. The initial condition was counterbalanced across animals.

Electro-mechanical stimulation was applied by using a magnetic field from a solenoid coupling to iron filings attached to both antennae. Iron filings mixed with epoxy were applied to the antennae and allowed to dry overnight to enable magnetic coupling. The magnetic field was generated by a custom-built solenoid electromagnet actuator powered by a regulated 12 V DC supply and controlled via an Arduino microcontroller. A push-button trigger initiated a pulse train consisting of 10 pulses delivered over 15 s (0.67 Hz; 50% duty cycle, 0.75 s ON / 0.75 s OFF). Stimulation was turned on when the animal reached the end of the platform and was switched off 5 s before the presentation of the looming stimuli.

### Extracellular recordings

Extracellular recordings from the Ventral Nerve Cord (VNC) were obtained from adult *Hieroglyphus banian* using custom-made hook electrodes. Animals were placed ventral side up in a custom holder and immobilized using modeling clay and wax. The forelegs and midlegs were removed, and a window was cut in the abdominal cuticle to expose the VNC. Connectives between the mesothoracic and metathoracic ganglia were isolated by removing fat bodies and tracheae. Extracellular DCMD recordings were made using a 280 µm insulated copper wire hook electrode (insulation removed at the contact site) positioned with a micromanipulator (MP-225, Sutter Instruments) on the ventral nerve cord between the mesothoracic and metathoracic ganglia. A chlorided silver wire inserted in the abdomen served as ground. Signals were amplified (200X; Axoclamp 900A, Molecular Devices), band-pass filtered (300 Hz–4 kHz; first-order high-pass, fourth-order Bessel low-pass), and digitized at 10 kHz (Digidata 1440A, Molecular Devices) using Clampex 10.3. Preparations were continuously perfused with insect Ringer’s solution (in mM: 140 NaCl, 5 CaCl_2_, 5 KCl, 4 NaHCO_3_, 6.3 HEPES, 1 MgCl_2_; pH 7.1). Visual stimuli were presented such that one compound eye was aligned to the center of the display.

### Data analysis and spike statistics

Spike times were extracted using custom Python scripts (NumPy, SciPy, Matplotlib). Spikes were binned at *Δt* = 1/60 *s*. Let *C*_*k,i*_ denote the spike count in trial *k* and bin *i*. Spike rate was computed as *R*_*k,i*_ = *C*_*k,i*_ / *Δt*: If there were N trials, the trial-averaged mean spike rate of *i*^*th*^ the bin was calculated as 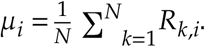. The standard deviation across trials was 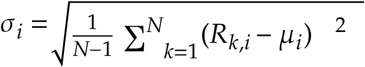. The standard error of the mean (SEM) was: 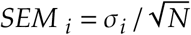. PSTHs were smoothed using a moving average filter of length for visualization purposes only. The trial-to-trial variability relative to the mean spike count was calculated using Fano factor: 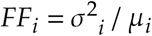.

### Spike rate comparisons and gain analysis

For each trial, spike counts within a 300 ms window around peak response were divided by window duration to obtain the mean spike rate. For each animal, rates were averaged across trials to yield one value per condition. Paired comparisons between conditions were performed using a paired *t*-test. Point-wise comparisons between conditions were computed as: 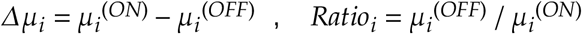. These measures quantify additive and multiplicative modulation of firing rate.

### Statistical analysis

Binary behavioral outcomes (jump vs no jump; grooming vs no grooming) were compared using Generalized Estimating Equations (GEE). Sample sizes (N), statistical tests, and exact p-values are reported in the corresponding figure legends or main text.

## STAR★METHODS

### Key resouces table

Key resources table
| REAGENT OR RESOURCE | SOURCE | IDENTIFIER |
| --- | --- | --- |
| **Experimental models: Organisms/strains** |  |  |
| *Hieroglyphus banian* (adult male and female) | Laboratory colony, Centre for Neural and Cognitive Sciences, University of Hyderabad | N/A |
| **Chemicals, peptides, and recombinant proteins** |  |  |
| Sodium chloride (NaCl) | Hi-Media (MB 023) | https://www.sigmaaldrich.com |
| Calcium chloride (CaCl<sub>2</sub>) | Hi-Media (TC 097) | https://www.sigmaaldrich.com |
| Potassium chloride (KCl) | Sigma-Aldrich (P 9541) | https://www.sigmaaldrich.com |
| Sodium bicarbonate (NaHCO<sub>3</sub>) | Sigma-Aldrich (S 5761) | https://www.sigmaaldrich.com |
| HEPES | SRL (68744) | https://www.sigmaaldrich.com |
| Magnesium chloride (MgCl<sub>2</sub>) | Sigma-Aldrich (M 1028) | https://www.sigmaaldrich.com |
| **Software and algorithms** |  |  |
| Python | Python Software Foundation 3.10.12 | https://www.python.org |
| Inkscape | Inkscape Project v1.4 | https://inkscape.org |
| Kdenlive | KDE Community v24.05.2 | https://kdenlive.org |
| **Other** |  |  |
| Axoclamp 900A amplifier | Molecular Devices | Model 900A |
| Digidata 1440A digitizer | Molecular Devices | Model 1440 |

