## Supplementary text and data for "Failures that are not failures: unobserved sensory history predicts escape behavior"

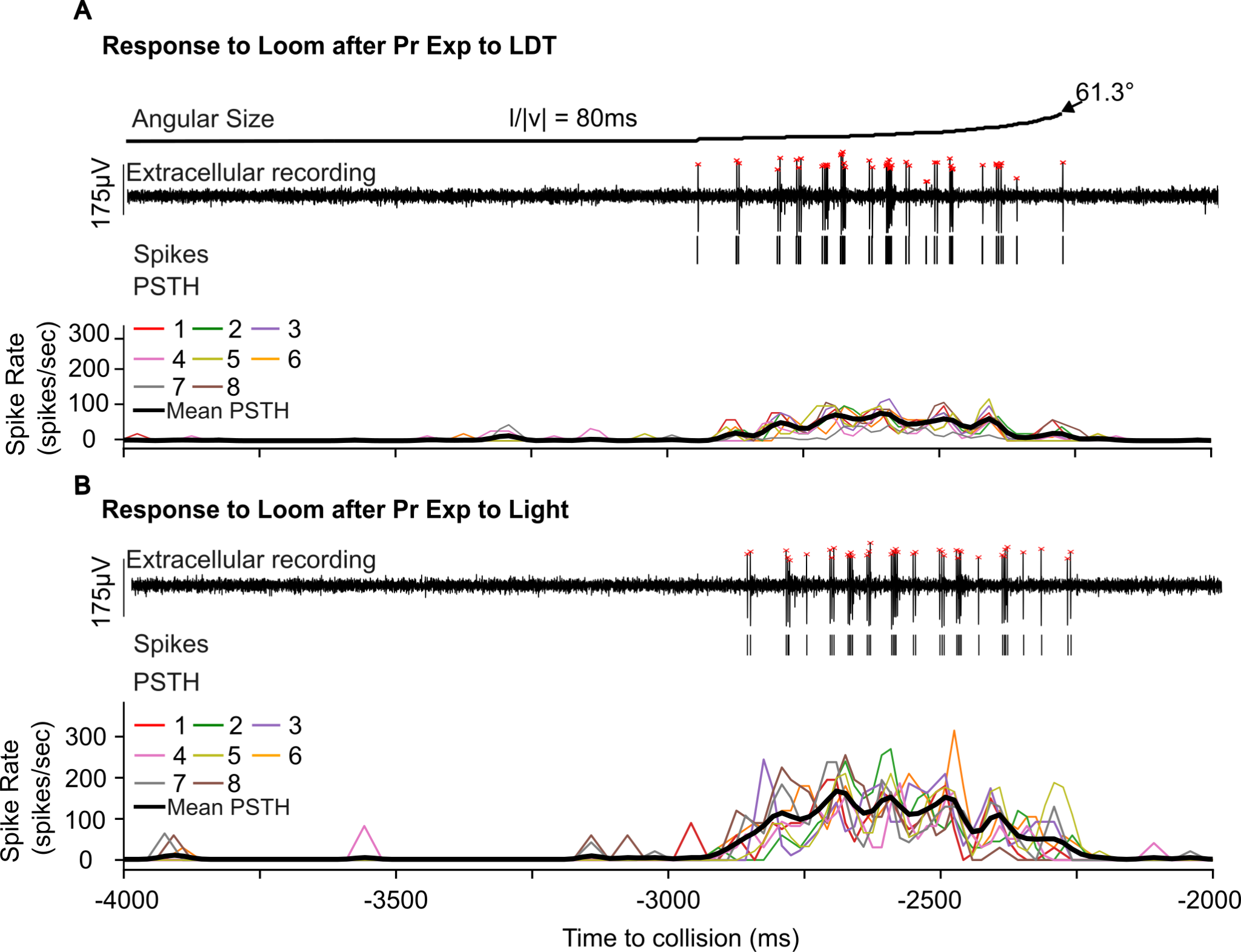

**Figure S1. Representative DCMD responses to looming stimuli under different light pre-exposure conditions.
(A)** Response to looming stimuli following pre-exposure to LDT. The top panel shows the looming stimulus profile, with angular size increasing over time (l/|v| = 80 ms) to a maximum of ~61.3°. The middle panel shows a representative extracellular recording of DCMD activity during looming stimulus presentation, with detected spikes indicated by red markers. The bottom panel shows peri-stimulus time histograms (PSTHs) for individual trials (colored traces, trials 1–8) and their mean (black trace).

**(B)** Same as (A) but response following pre-exposure to ‘light’ condition. PSTHs were computed using a bin width of 1/60 s and smoothed with a rectangular moving average filter (window length = 4 bins).

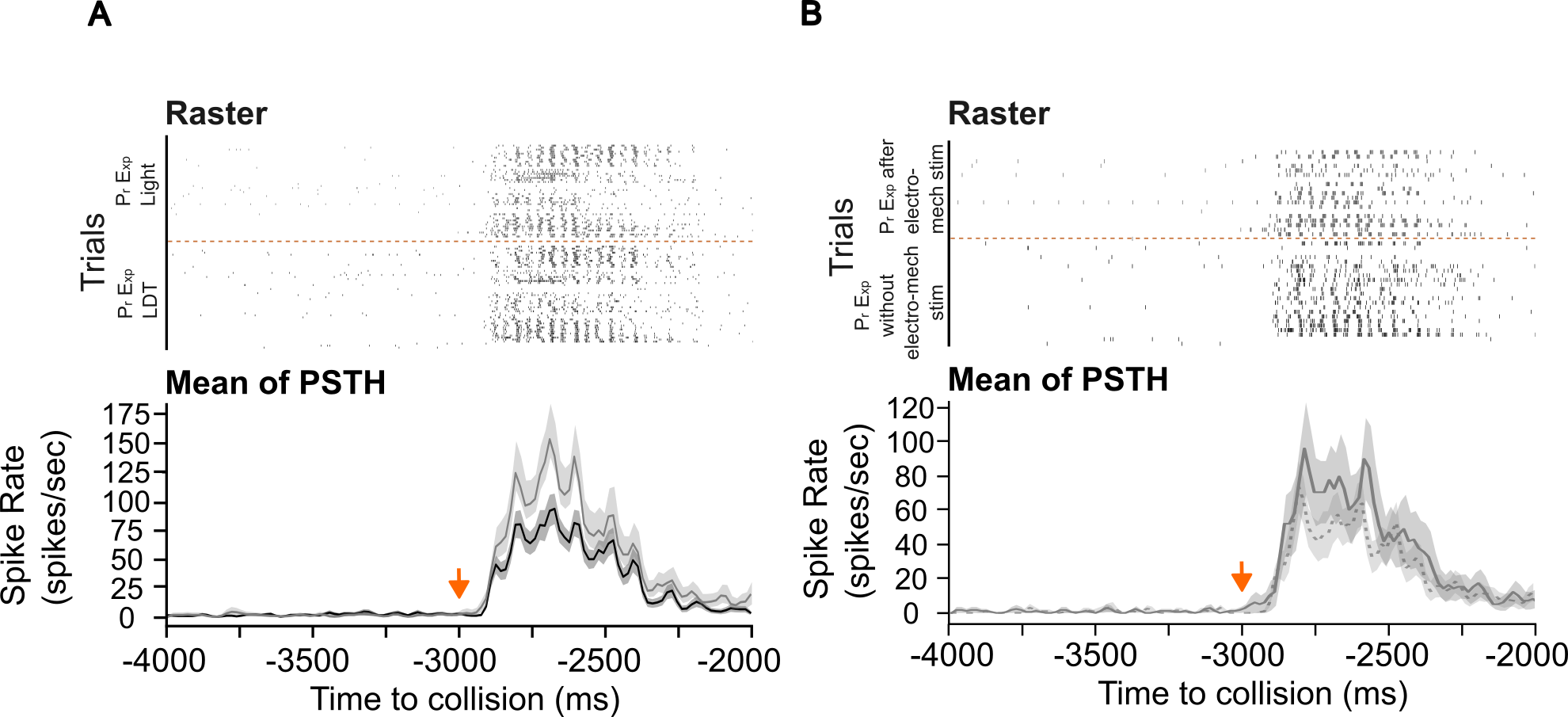

**Figure S2: Raster plots and mean PSTHs of looming-evoked DCMD responses under different pre-treatments.**

**(A)** Raster plots showing DCMD firing across all trials collected from 7 animals in response to looming stimuli, comparing LDT and Light pre-exposures. Each row represents one trial, aligned to time to collision. Trials from the two pre-exposure conditions are separated by a dotted line. The lower panel shows the corresponding mean PSTHs across animals for each condition, plotted as mean ± SEM (black: loom response after LDT pre-exposure; grey: loom response after Light pre-exposure).

**(B)** Same as (A), but for recordings from 5 animals comparing looming-evoked responses following electro-mechanical pre-exposure that induces grooming (above dotted line) and no electro-mechanical stimulation pre-exposure (below dotted line). Corresponding mean of PSTHs across animals are shown in the lower panel (mean ± SEM).

All PSTHs were computed using a bin width of 1/60 s. Average PSTHs were smoothened using a rectangular moving-average filter (window length = 4 bins). The orange arrow indicates looming stimuli onset.

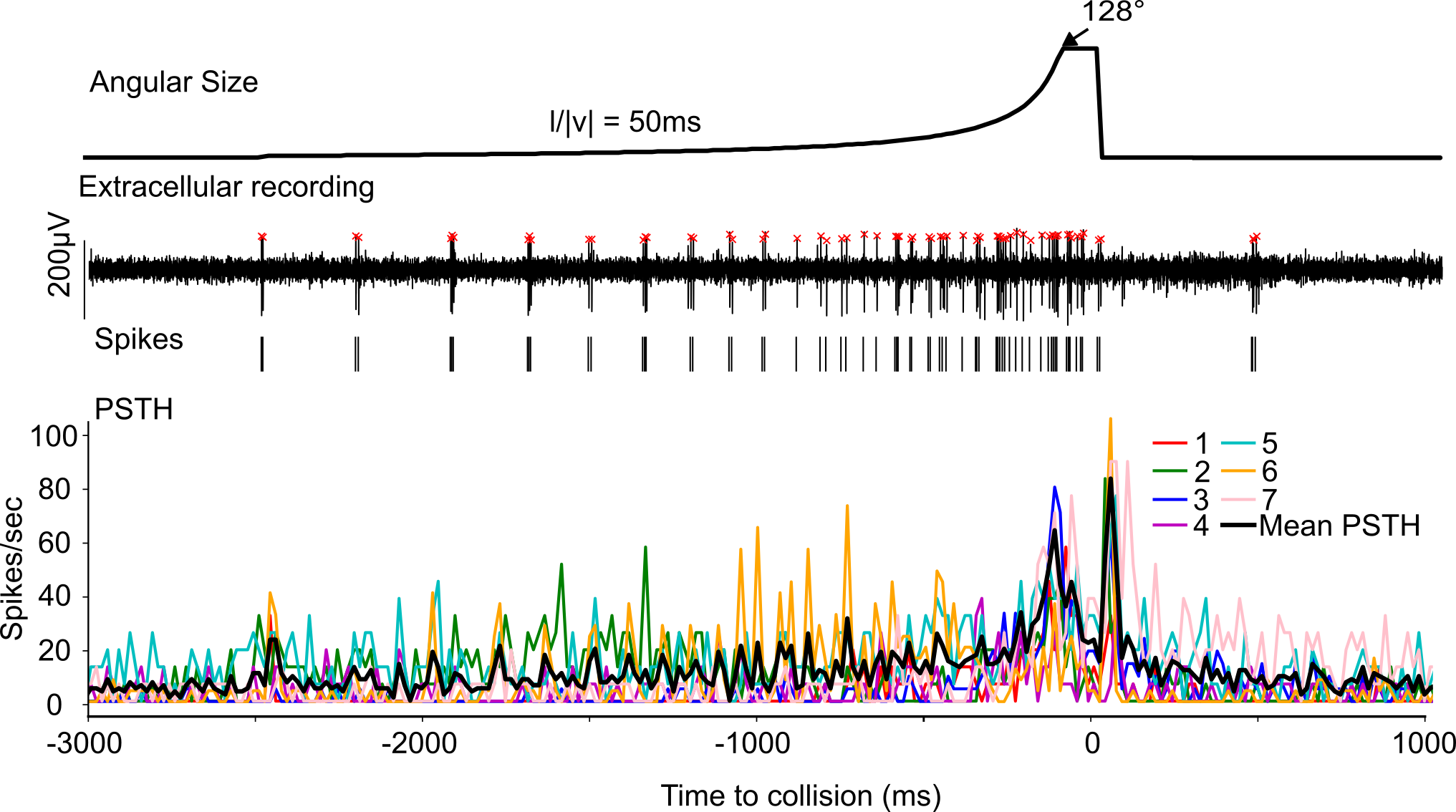

**Figure S3: Representative looming-evoked DCMD activity under an l/|v| = 50 ms stimulus condition.**

The top panel shows the looming stimulus profile, with angular size increasing over time (l/|v| = 50 ms) to a maximum of ~128°. The middle panel shows a representative extracellular recording of DCMD activity during looming stimuli presentation, with detected spikes indicated by red markers. The bottom panel shows PSTHs for individual trials (colored traces, trials 1–7) and their mean (black trace), computed using a bin width of 1/60 s and smoothened with a rectangular moving-average filter (window length = 4 bins).

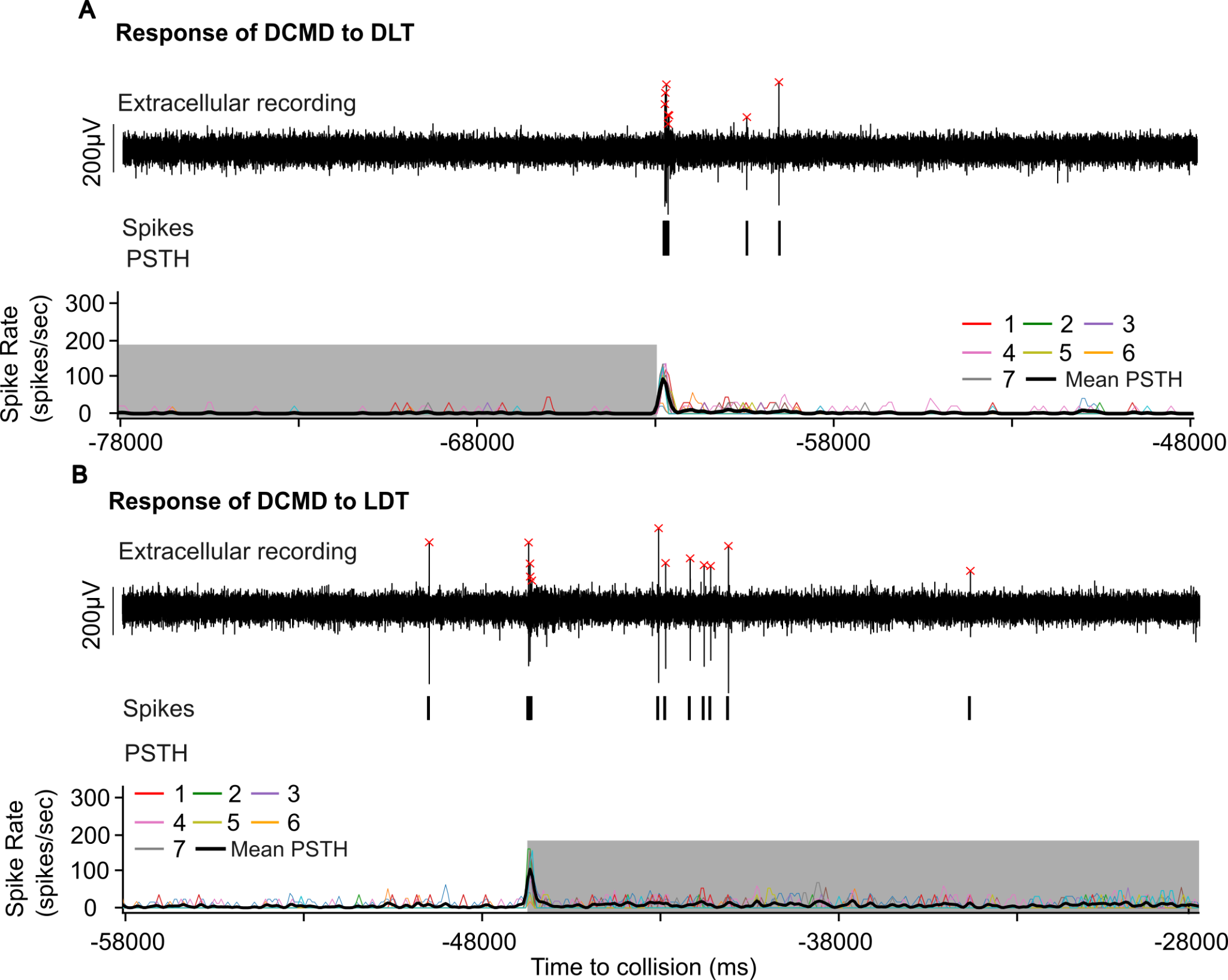

**Figure S4: Representative DCMD responses to DLT (Dark to light transition) and LDT (Light to dark transition).**

**(A)** Response to DLT. The top panel shows a representative extracellular recording of DCMD activity during DLT, with detected spikes indicated by red markers. The bottom panel shows peri-stimulus time histograms (PSTHs) for individual trials (colored traces, trials 1–7) and their mean (black trace).

**(B)** Same as (A) but response to LDT. PSTHs were computed using a bin width of 1/60 s and smoothed with a rectangular moving average filter (window length = 4 bins).

The dark gray and light gray shaded regions indicate dark and light periods respectively.

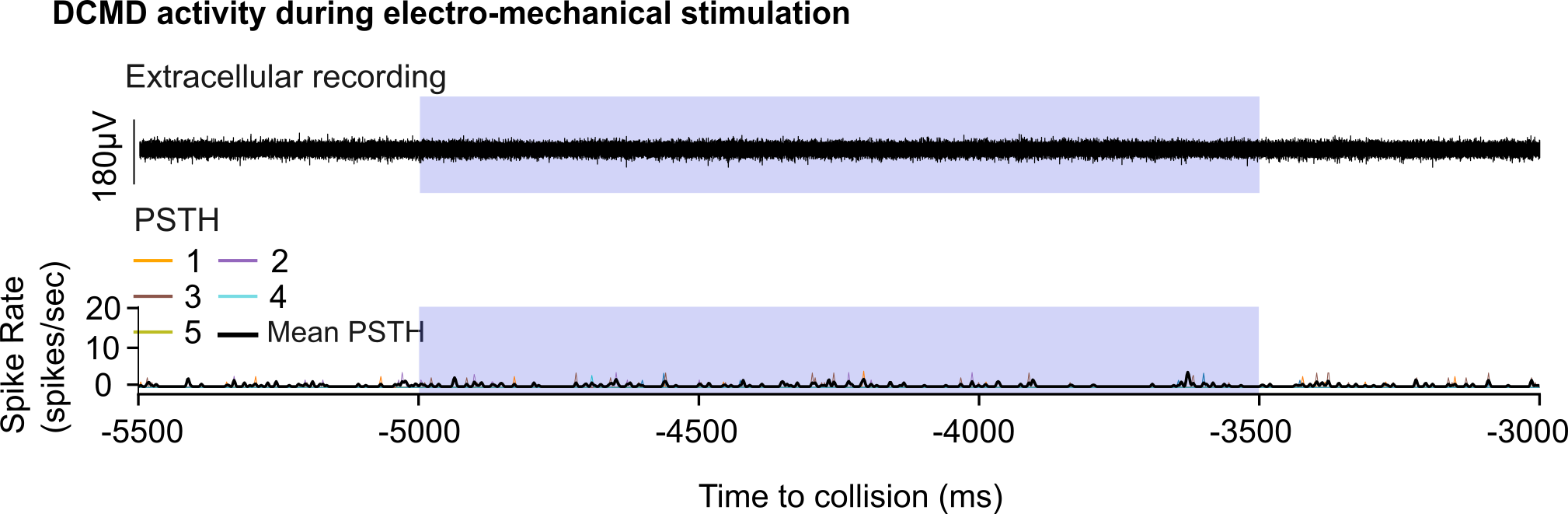

**Figure S5: Representative DCMD responses to electro-mechanical stimulation.**

The top panel shows a representative extracellular recording of DCMD activity during electro-mechanical stimulation. The bottom panel shows PSTHs for individual trials (colored traces, trials 1–5) and their mean (black trace), computed using a bin width of 1/60 s and smoothened with a rectangular moving-average filter (window length = 4 bins). The blue shaded region indicates the electro-mechanical stimulation period.

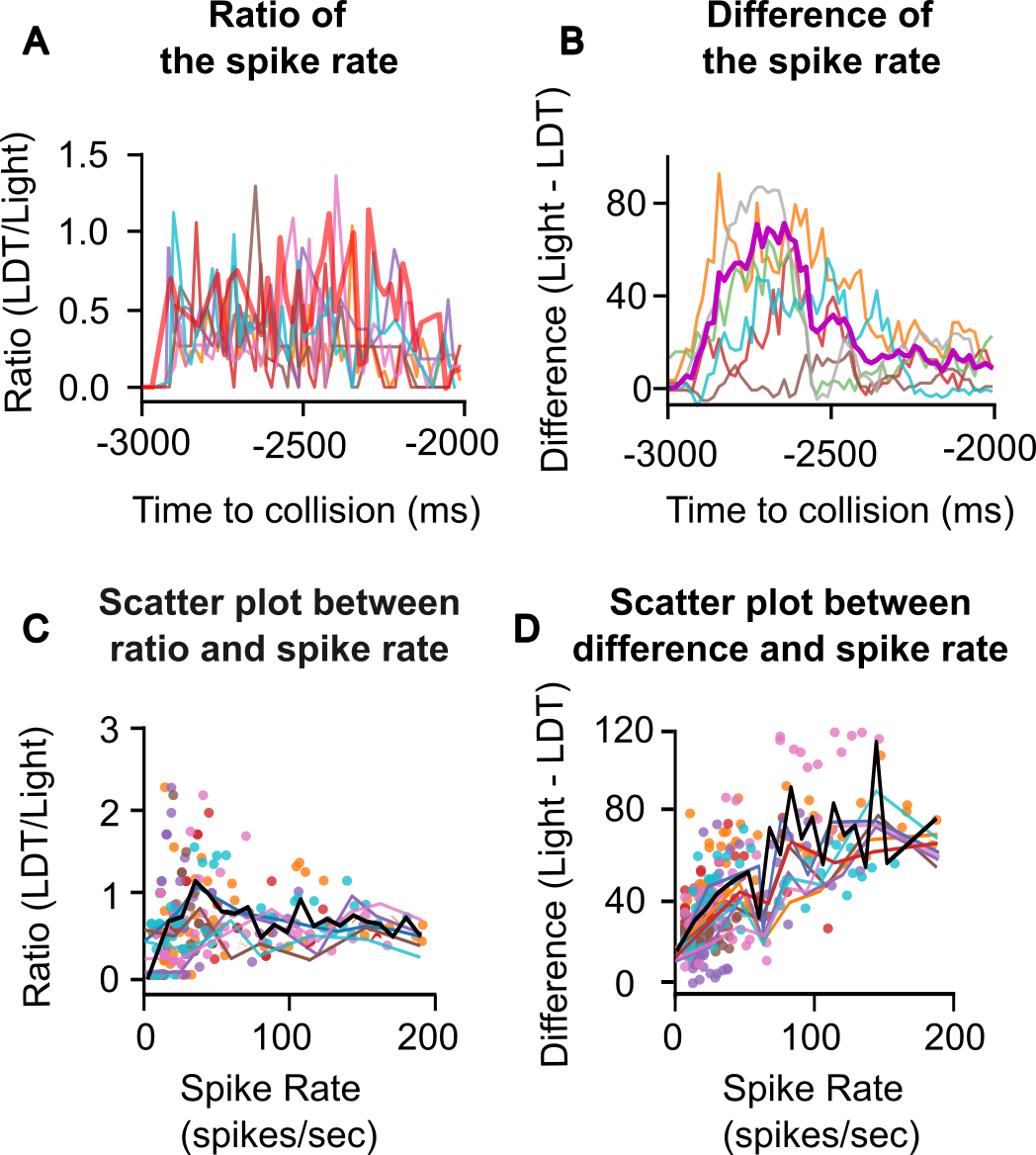

**Figure S6: Relationship of the difference and ratio of DCMD firing rate to mean firing rates, demonstrating divisive gain modulation following LDT pre-exposures.**

**(A)** Ratio of DCMD firing rate (LDT / Light) during the looming response for individual animals (colored traces) and the population mean (red). The ratio was calculated from the trial-averaged PSTH of each animal (PSTH bin width = 1/60 s).

**(B)** Difference in firing rate (Light − LDT) during the looming response for individual animals (colored traces) and the population mean (purple).

**(C)** Scatter plot showing the relationship between firing rate (mean response across Light and LDT pre-exposure conditions) and the firing rate ratio (LDT / Light). Each point represents one PSTH time bin from an individual animal. Colored lines represent the binned mean for each animal, and the black line represents the population mean.

**(D)** Scatter plot showing the relationship between firing rate and the difference in firing rates (Light − LDT). Each point represents one PSTH time bin from an individual animal. Colored lines indicate the binned mean for each animal, and the black line represents the population mean.

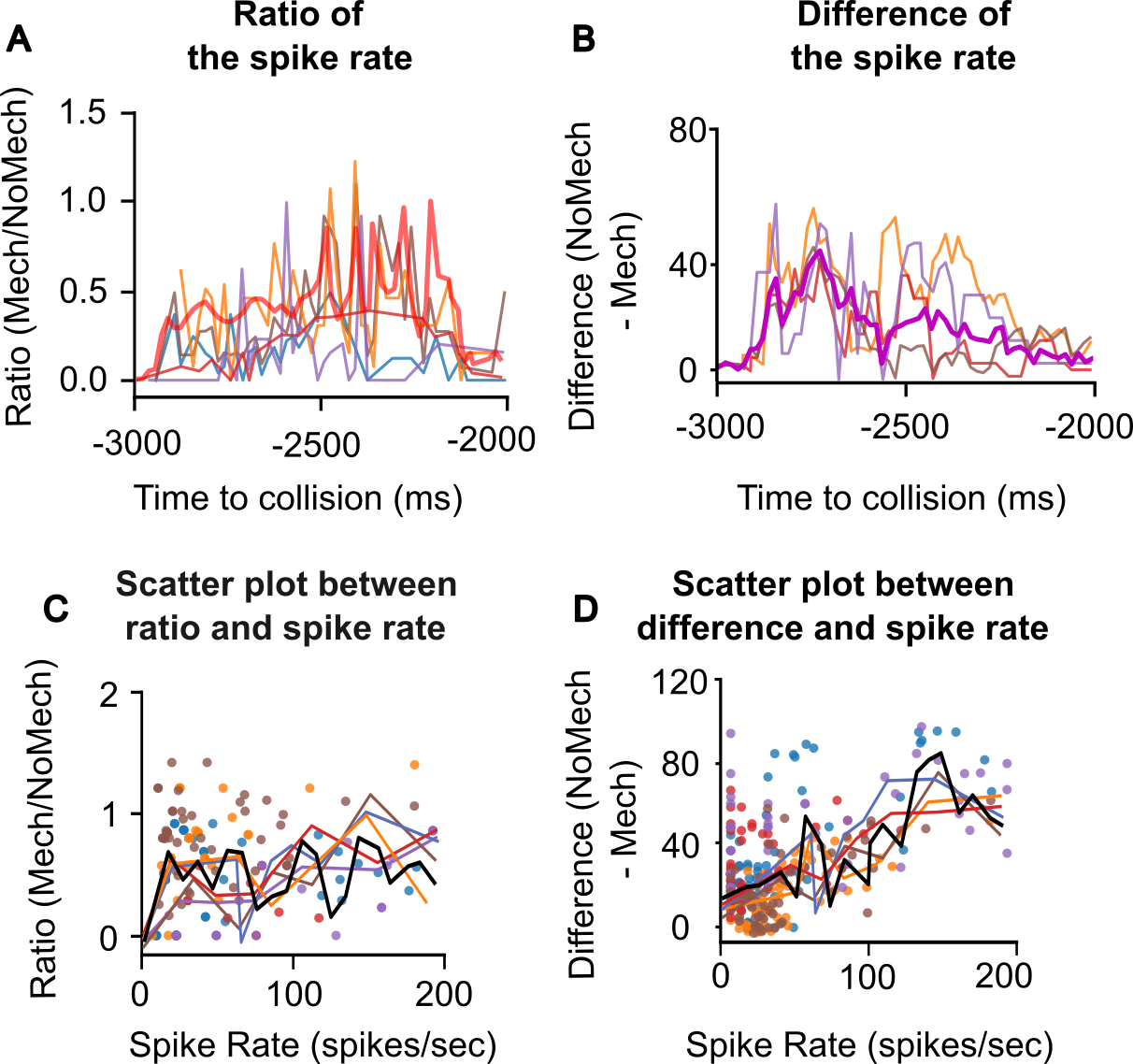

**Figure S7. Relationship of the difference and ratio of DCMD firing rates to mean firing rate, demonstrating divisive gain modulation following electromechanical (Mech) stimulation.**

**(A)** Ratio of DCMD firing rate (Mech / NoMech) during the looming response for individual animals (colored traces) and the population mean (red). The ratio was calculated from the trial-averaged PSTH of each animal (PSTH bin width = 1/60 s).

**(B)** Difference in firing rate (NoMech − Mech) during the looming response for individual animals (colored traces) and the population mean (purple).

**(C)** Scatter plot showing the relationship between firing rate (mean response across Mech and NoMech conditions) and the firing rate ratio (Mech / NoMech). Each point represents one PSTH time bin from an individual animal. Colored lines represent the binned mean for each animal, and the black line represents the population mean.

**(D)** Scatter plot showing the relationship between firing rate and the difference in firing rate (NoMech − Mech). Each point represents one PSTH time bin from an individual animal. Colored lines indicate the binned mean for each animal, and the black line represents the population mean.

**Supplementary Movies:**

**Movie S1:** Example video recording of a trial in which the grasshopper jumped in response to a looming stimulus. ($l/|v|$ = 80 ms). The video is shown at 50% of original speed (60fps).

**Movie S2:** Example video of a grasshopper showing grooming behavior in the culture chamber during the dark phase. The video is shown at 50% of original speed (60fps).
